# Specialized actin nanoscale layers control focal adhesion turnover

**DOI:** 10.1101/2023.02.15.528622

**Authors:** Reena Kumari, Katharina Ven, Megan Chastney, Johan Peränen, Jesse Aaron, Leonardo Almeida-Souza, Elena Kremneva, Renaud Poincloux, Teng-Leong Chew, Peter W. Gunning, Johanna Ivaska, Pekka Lappalainen

**Author notes:** Corresponding author: Pekka Lappalainen, HiLIFE Institute of Biotechnology, P.O. Box 56, 00014 University of Helsinki, Finland.

## Abstract

Focal adhesions (FAs) connect inner workings of the cell to the extracellular matrix to control cell adhesion, migration, and mechanosensing^1,2^. Previous studies demonstrated that FAs contain three vertical layers, which connect extracellular matrix to the cytoskeleton^3,4,5^. However, cellular processes rely on precisely-regulated FA turnover, but the molecular machineries that control FA assembly and disassembly have remained elusive. By using super-resolution iPALM microscopy, we identified two unprecedented nanoscale layers within FAs, specified by actin filaments bound to tropomyosin isoforms Tpm1.6 and Tpm3.2. The Tpm1.6-actin filaments beneath the previously identified ‘actin-regulatory layer’ are critical for adhesion maturation and controlled cell motility, whereas the Tpm3.2-actin filament layer towards the bottom of FA facilitates adhesion disassembly. Mechanistically, Tpm3.2 stabilizes KANK-family proteins at adhesions, and hence targets microtubule plus-ends to FAs to catalyse their disassembly. Loss of Tpm3.2 leads to disorganized microtubule network, abnormally stable FAs, and defects in tail retraction during cell migration. Thus, FAs are composed of at least three distinct actin filament layers, each having specific roles in coupling of adhesion to the cytoskeleton, or in controlling adhesion dynamics. In a broader context, these findings demonstrate how distinct actin filament populations can co-exist and perform specific functions within a defined cellular compartment.

## MAIN

Interplay between cells and extracellular matrix (ECM) is critical for a range of physiological and pathological processes. For example, in embryonic development and during immune responses in adult tissues, cells must sense both chemical and mechanical properties of their environment for decision-making in proliferation, migration, and differentiation. The most prominent structures that link animal cells to ECM are focal adhesions (FAs). These complex protein assemblies interact with ECM through transmembrane integrin adhesion receptors, which are then connected to the actin cytoskeleton through a large number of associated proteins. Defects in the architecture or regulation of FAs are associated with several pathological states, such as cancer metastasis and immunological disorders ^1,2^.

At the leading edge of migrating mesenchymal cells, small (diameter <250 nm) *nascent adhesions* form within the actin-rich lamellipodia. A subset of *nascent adhesions* undergoes maturation to *focal complexes* at the lamellipodium-lamella interface, and can further mature to *focal adhesions* in response to increased contractile forces from the associated actomyosin bundles, called stress fibers, or through externally applied forces to the cell ^6,7,2^. Elevated traction forces also inhibit elongation of the FA–associated stress fibers to further elevate the contractile force exposed to the adhesion ^8^. Hence, FA assembly and maturation are tightly regulated by the organization of the actin cytoskeleton. However, if the actin cytoskeleton also contributes to other aspects of adhesion dynamics remains poorly defined.

In addition to assembly, the disassembly of FAs must also be tightly regulated. This is especially important during directional cell migration, where adhesion disassembly occurs towards the cell rear to allow tail retraction for movement ^9^. The precise molecular mechanisms of adhesion disassembly are incompletely understood, but were shown to involve endocytosis of adhesion-resident integrins and local proteolysis of ECM components ^10, 2^. Moreover, targeting microtubules to FAs has for long been known to trigger adhesion disassembly ^11–14^. KANK family proteins, which interact with talin through their Kank amino-terminal (KN) domain, were identified as critical factors for microtubule-dependent disassembly of adhesions ^15, 16^. At the molecular level, KANK-dependent association of microtubule plus-ends with FAs promotes adhesion turnover, at least partly by trapping RhoA GEF-H1 to microtubules, and hence inhibiting the local activation of RhoA ^17^. Moreover, microtubules were shown to deliver autophagosomes to mature FAs to promote their disassembly ^18–20^. Therefore, the spatio-temporal interplay between microtubules and FAs is a complex process, which is also likely to involve other, currently unidentified components. Indeed, proteomics studies provided evidence that adhesions harbour hundreds of different proteins ^21–25^, but the functions of only fraction of these have been identified so far.

A major breakthrough in understanding FA architecture came with the use of super-resolution interferometric photo-activated localization microscopy (iPALM), revealing that FAs are composed of at least three separate layers relative to the plasma membrane: 1). ‘Integrin signalling layer’ at the bottom of adhesion composed of transmembrane integrins, FAK, and paxillin, 2). ‘Force-transduction layer’, composed of vinculin and extended talin molecules, which link integrins to actin filaments, and 3). Uppermost ‘Actin-regulatory layer’, containing α-actinin cross-linked actin filaments and actin polymerizing factor VASP ^3–5, 26^. However, actin filaments within adhesions display wider vertical distribution compared to that of α-actinin, hinting at the presence of other, currently unidentified actin filament structures in FAs ^27^. Moreover, in addition to VASP, several other actin filament assembly factors were shown to associate with focal adhesions ^20, 28–30^. Thus, the presence of a large number of actin-regulatory and signalling proteins, together with the need to precisely control FA assembly, maturation and disassembly during various cellular processes, proposes that the molecular organization of FAs may be more complex than depicted in current models.

### Tpm1.6 and Tpm3.2 generate specific actin filament layers along the vertical axis of focal adhesions

A key feature of actin filaments is their ability to assemble into functionally distinct arrays in cells ^31–35^. Tropomyosins (Tpms) are ubiquitous actin-binding proteins, which have evolved to specify the functional diversity of cellular linear actin filament arrays ^36, 37^. The coiled-coil tropomyosin dimers form head-to-tail polymers along the major groove of actin filaments, and are only known to be functional when bound to actin filaments ^38, 39^. The four *tropomyosin* genes in mammals, *TPM1*, *TPM2*, *TPM3* and *TPM4*, can generate >40 Tpm isoforms through alternative splicing, and the different isoforms provide specific functional features to the associated actin filaments ^40–43^. Several Tpm isoforms localize to actin stress fibers and regulate their assembly. From these, at least two isoforms, Tpm1.6 and Tpm3.2, appear to also co-localize with focal adhesions ^44^. Importantly, Tpm1.6 and Tpm3.2 cannot co-polymerize with each other on the same actin filament. Moreover, both isoforms compete with α-actinin, the key component of the previously identified ‘actin-regulatory layer’ of focal adhesions, for actin filament binding ^41, 45, 46^. Thus, if Tpm1.6 and Tpm3.2 are indeed focal adhesion components, they may specify previously unknown, functionally distinct actin filament structures.

To investigate Tpm1.6 and Tpm3.2 localization in focal adhesions, we employed total internal reflection fluorescence microscopy (TIRFM) imaging to detect structures localizing immediately adjacent to the cell-ECM interface. Using specific antibodies ^47^ and fluorescent fusion proteins, we detected Tpm1.6 and Tpm3.2 localizing to FAs (Fig. 1a; S1a), in agreement with a previous studies using wide-field microscopy ^44^. Tpm1.6 was distributed across the entire length of the adhesion and extended to the associated dorsal stress fiber, whereas Tpm3.2 localization was predominantly restricted to the adhesion (Fig. 1a-c, Fig. S1b). Both tropomyosins partially co-localized with α-actinin-1 in adhesion (Fig. S1c). Please note that the pronounced localization of α-actinin-1 to the distal ends of focal adhesions is most likely due to enrichment of α-actinin-1 at the lamellipodial actin network ^48^. Fluorescence-recovery-after-photobleaching (FRAP) analysis demonstrated that both tropomyosins are relatively stable components of FAs. However, consistent with earlier *in vitro* work on purified proteins ^41^, Tpm3.2 displayed more rapid association-dissociation dynamics in FAs compared to Tpm1.6 (Fig. S1d-e). Moreover, TIRF live-cell imaging on U2OS cells expressing EGFP-Tpm1.6, mRuby2-Tpm3.2, and miRFP670-Paxillin revealed that Tpm1.6 and Tpm3.2 were recruited to the newly forming adhesions simultaneously with paxillin. However, both Tpm1.6 and Tpm3.2 intensities increased rapidly over time, whereas paxillin intensity gradually increased at adhesions, (Fig. 1d-f, Supplementary movie 1). Thus, Tpm1.6 and Tpm3.2 are early components of focal adhesions that display different dynamics and slightly different lateral (xy) localization patterns with each other, and with α-actinin, in adhesions.

**Figure 1.**
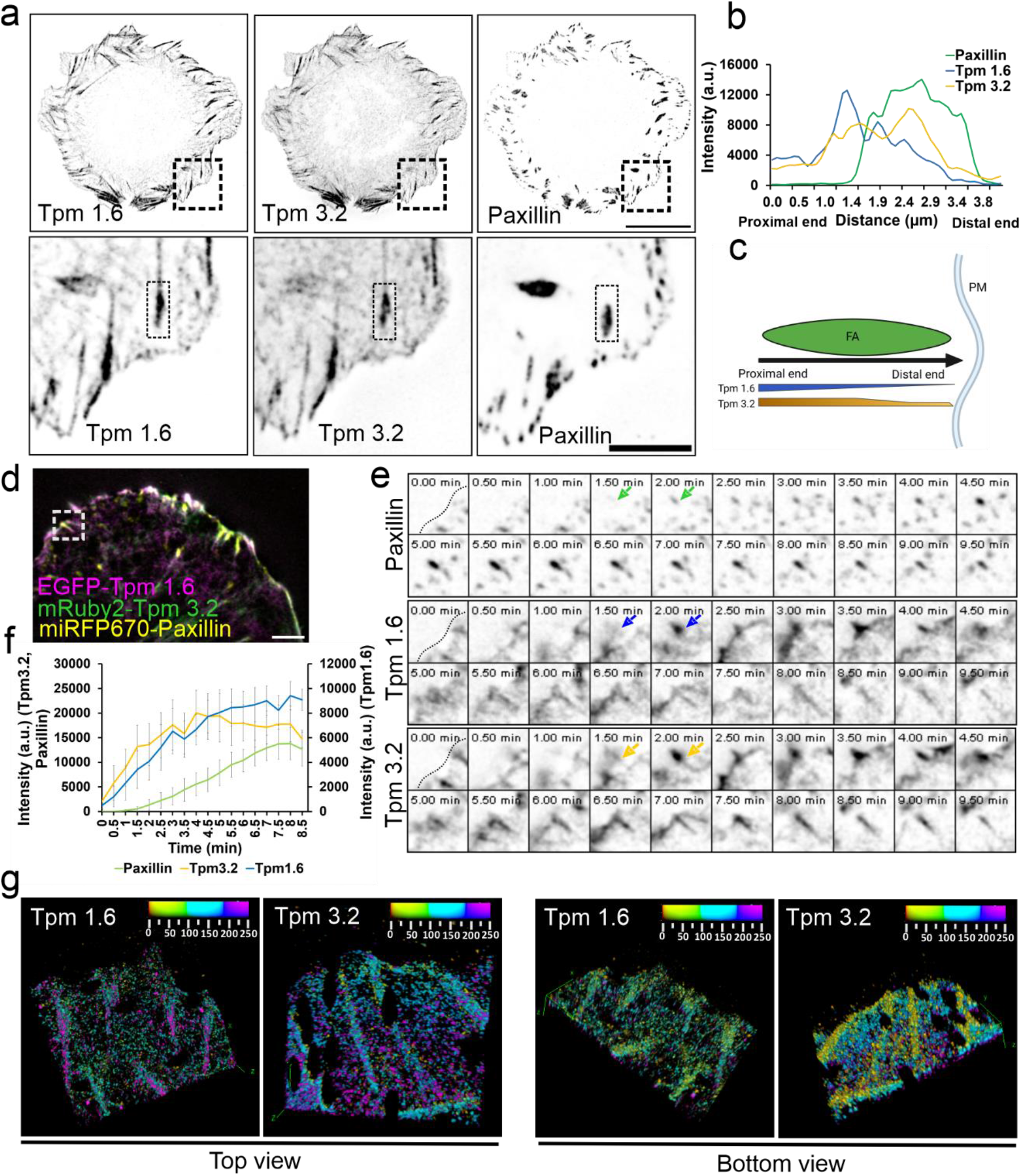
Tropomyosin-1.6 and tropomyosin-3.2 are early components of focal adhesions. (a) Representative TIRF images of wild-type U2OS cells expressing pmRuby2C1-Tpm1.6 and pEGFPC1-Tpm3.2, and stained for endogenous paxillin. The panels at the bottom are magnified images of the regions indicated with dashed boxes highlighting the localization of Tpm1.6 and Tpm3.2 in focal adhesions. Scale bars, 20 μm and 5 μm in upper and bottom rows, respectively. (b) Representative line scan intensity profiles across the focal adhesion (from panel ‘a’). (c) Schematic representation of lateral localizations of Tpm1.6 and Tpm3.2 from the proximal to distal end of focal adhesions (see also Fig. S1b). (d) TIRF image from a time-lapse movie of a wild-type U2OS expressing miRFP670-Paxillin, pEGFPC1-Tpm1.6 and pmRuby2C1-Tpm3.2. Scale bar, 50 μm. (e) Individual channels of selected time-lapse frames from the magnified region (indicated by a white box in panel ‘d’). Black dotted lines indicate the cell edge. Green, purple and yellow arrows highlight the timing of paxillin, Tpm1.6 and Tpm3.2 accumulation to the focal adhesion, respectively. (f) Intensity profile analysis (n=5 adhesions) related to panel d and e, demonstrating that Tpm1.6 and Tpm3.2 become enriched at focal adhesions before the recruitment of paxillin. The graph represents mean ± SE (g) 3D-top view (image orientation from dorsal plane) and 3D-bottom view (image orientation from ventral plane) of rendered iPALM images of Tpm1.6 and Tpm3.2. The images were obtained by using ‘3D viewer’ plugin from FIJI/ImageJ. Note the enrichment of Tpm1.6 at the dorsal side (in top-view) and enrichment of Tpm3.2 at the ventral side (in bottom-view).

To examine the nanoscale localisation of Tpm1.6- and Tpm3.2-actin filaments in FAs, we applied iPALM, which allows 3D localization of tagged proteins with <20 nm accuracy ^3–5, 26, 27^. First, we generated Eos-tagged Tpm1.6 and Tpm3.2 and validated their localization to FAs of U2OS cells together with α-actinin by TIRF imaging (Fig. S1f). Interestingly, subsequent iPALM experiments provided evidence that Tpm1.6 and Tpm3.2 display distinct vertical localizations in respect to each other (Fig. 1g; Fig. S2a,b; Supplementary movie 2). To dissect the precise localization of Tpm1.6 and Tpm3.2 in the nanoscale strata of FAs, we used endogenous paxillin (visualized by antibody) as a reference marker for focal adhesions and the ‘integrin signalling layer’, and established photoactivatable fluorescent (PA-FP) proteins of talin-N and α-actinin-1 ^3, 27^ as reference proteins for the ‘force-transduction layer’ and ‘actin-regulatory layer’, respectively ^3–5, 27^. The vertical (*z*) localization of paxillin (Z_centre_ = 51 nm ± 14 nm) was consistent with its previously reported z-position in U2OS and other cell-types (Fig. 2a, f). The observed z-positions of talin-N and α-actinin were consistent with the previously reported z-positions in cornerstone adhesions of pluripotent stem cells ^27^, while slightly higher compared to the previous studies on U2OS and endothelial cells (see ‘Methods’ and Supplementary Table 1). The small variations may result from differences in the analysed adhesion types, or from the differences in coverslip coating for imaging.

**Figure 2.**
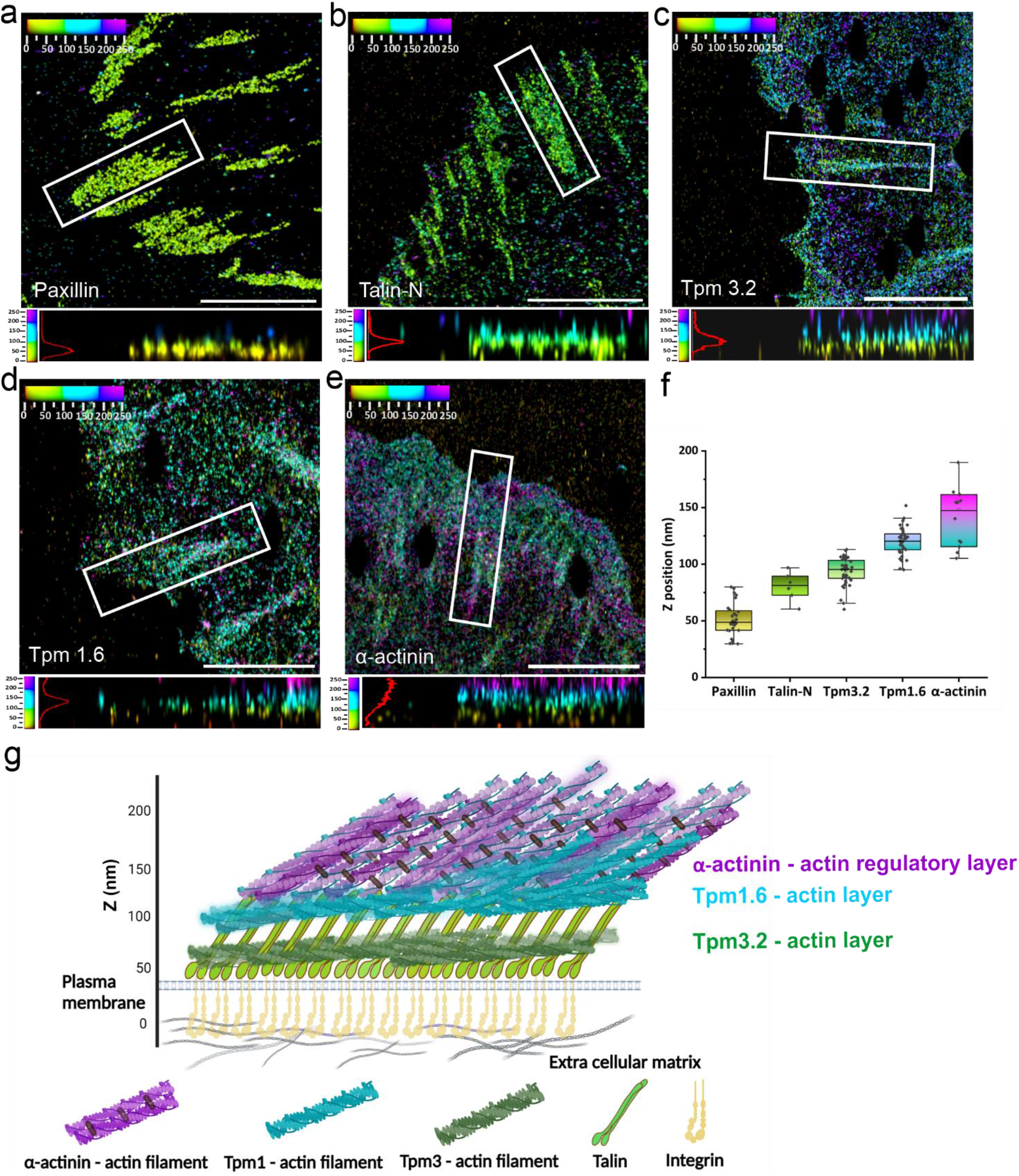
Tropomyosin-1.6 and tropomyosin-3.2 actin filament arrays form specific nanoscale layers in focal adhesions. (a-e) XY views and side views (indicated by white boxes in XY-view panels) form iPALM images of focal adhesions of U2OS cells. (a). Endogenous paxillin, (b) tdEos-Talin-N, (c) mEos3.2-Tpm3.2, (d) mEos3.2-Tpm1.6, and (e) mEos3.2-α-actinin. Scale bars, 5 μm. (f) Vertical stratification of focal adhesions located at the leading edge of the cell showing the Z-positions (Z_center_) of paxillin, Talin-N, Tpm3.2, Tpm1.6 and α-actinin. Each point in the graph corresponds to an individual focal adhesion measurement. Boxes display the mean, median, Whiskers, IQR: 25^th^-75^th^ percentiles, Whiskers range within 1.5*IQR. (g) Schematic model of the molecular 3D architecture of a focal adhesion displaying the localizations of three ‘actin filament layers’ in the nanoscale strata. The positioning of each protein is based on the data presented in panels ‘a-f’. The model does not depict the protein stoichiometry.

To focus on similar adhesion types, we first analysed only those adhesions, which are located at the leading edge and oriented perpendicular to the cell edge (Fig. S3a-e). Interestingly, our analysis revealed that the z-position of Tpm1.6 (Z_centre_ = 120 nm ± 12 nm) was slightly below the extended vertical distribution of α-actinin cross-linked actin filaments (Z_centre_ = 120 nm ± 15 nm to Z _centre_ = 175 nm ± 20 nm), and displayed only a partial overlap with α-actinin (‘actin-regulatory layer’) (Fig. 2d, e, f). Moreover, the overall distribution range of Tpm1.6 (spanning ~ 25 nm) did not correspond to the wider z-distribution of α-actinin (spanning ~ 50 nm). Strikingly, Tpm3.2 localized to markedly lower vertical position (Z_centre_ = 93 nm ± 12 nm) compared to α-actinin and Tpm1.6, showing partial overlap with talin-N (Z_centre_ = 80 nm ± 12 nm) in the ‘force transduction layer’ (Fig. 2b, c, f). Since the newly identified Tpm1.6- and Tpm3.2-actin arrays show distinct vertical nanoscale localizations in comparison to previously described ‘actin-regulatory layer’, we propose to call these newly identified actin layers as ‘Tpm1.6-actin filament layer’ and ‘Tpm3.2-actin filament layer’ (Fig. 2f). In addition to focal adhesions at the leading edge of cell, we analysed the vertical localizations of Tpm1.6 and Tpm3.2 in large, mature focal adhesions, which are associated with the ventral stress fibers and oriented more parallel along the cell edge. Also here, Tpm1.6 and Tpm3.2 localized to similar two distinct vertical layers, as described above for the leading edge adhesions (Fig. S2c-e).

Together, these live-cell and super-resolution iPALM imaging experiments reveal that focal adhesions are composed of at least three distinct actin-filament populations, which are specified by actin-binding proteins α-actinin, Tpm1.6 and Tpm3.2. Importantly, these actin filament populations display different localizations along the vertical axis of adhesions: The previously described ‘α-actinin-actin regulatory layer’ forming the upper-most layer, followed by partially overlapping ‘Tpm1.6-actin filament layer’, and finally a ‘Tpm3.2-actin filament layer’ towards the bottom of adhesion (Fig. 2g).

### Tpm1.6 and Tpm3.2 have different roles in cell morphogenesis and migration

Considering the distinct localizations of Tpm1.6 and Tpm3.2-actin filaments in the nanoscale strata of focal adhesions, we next determined the functions of these two actin-filament layers by generating knockout U2OS cell lines by CRISPR/Cas9. The obtained knockout clones were confirmed by Western blot, Sanger sequencing and Next Generation sequencing (Fig. S4a-d). Please note that by targeting exon 1a of the *TPM1* gene, and exon 1b of the *TPM3* gene, we only depleted Tpm1.6 and the closely related Tpm1.7, and Tpm3.2 and the closely related Tpm3.1, respectively, from the Tpm isoforms expressed in U2OS cells (Fig. S4a; ^49^). Depletions of Tpm1.6/7 (hereafter Tpm1 knockout cells) or Tpm3.1/2 (hereafter Tpm3 knockout cells) did not result in elevation or downregulation of the other isoform (Fig. S4b). Both Tpm1 and Tpm3 knockout cells displayed thinner and less organized actin stress fibers compared to the wild-type cells (Fig. 3a-b), in line with our previous siRNA studies ^44^. This stress fiber phenotype was confirmed by mixing the Tpm1 KO and Tpm3 KO cells with wild-type U2OS cells stably expressing EGFP-CAAX, and comparing the organization of actin networks in control and knockout cells from the same images (Fig. S4e). Consistent with the defects in the stress fiber networks, the Tpm1 and Tpm3 knockout cells generated diminished tractions forces (Fig 3d-e) and displayed slightly reduced levels of active RhoA (Fig. S4h). On the other hand, myosin II (both NMIIA and NMIIB) localization along stress fibers appeared largely unaffected in the Tpm1 and Tpm3 knockout cells (Fig. S4i-j).

**Figure 3.**
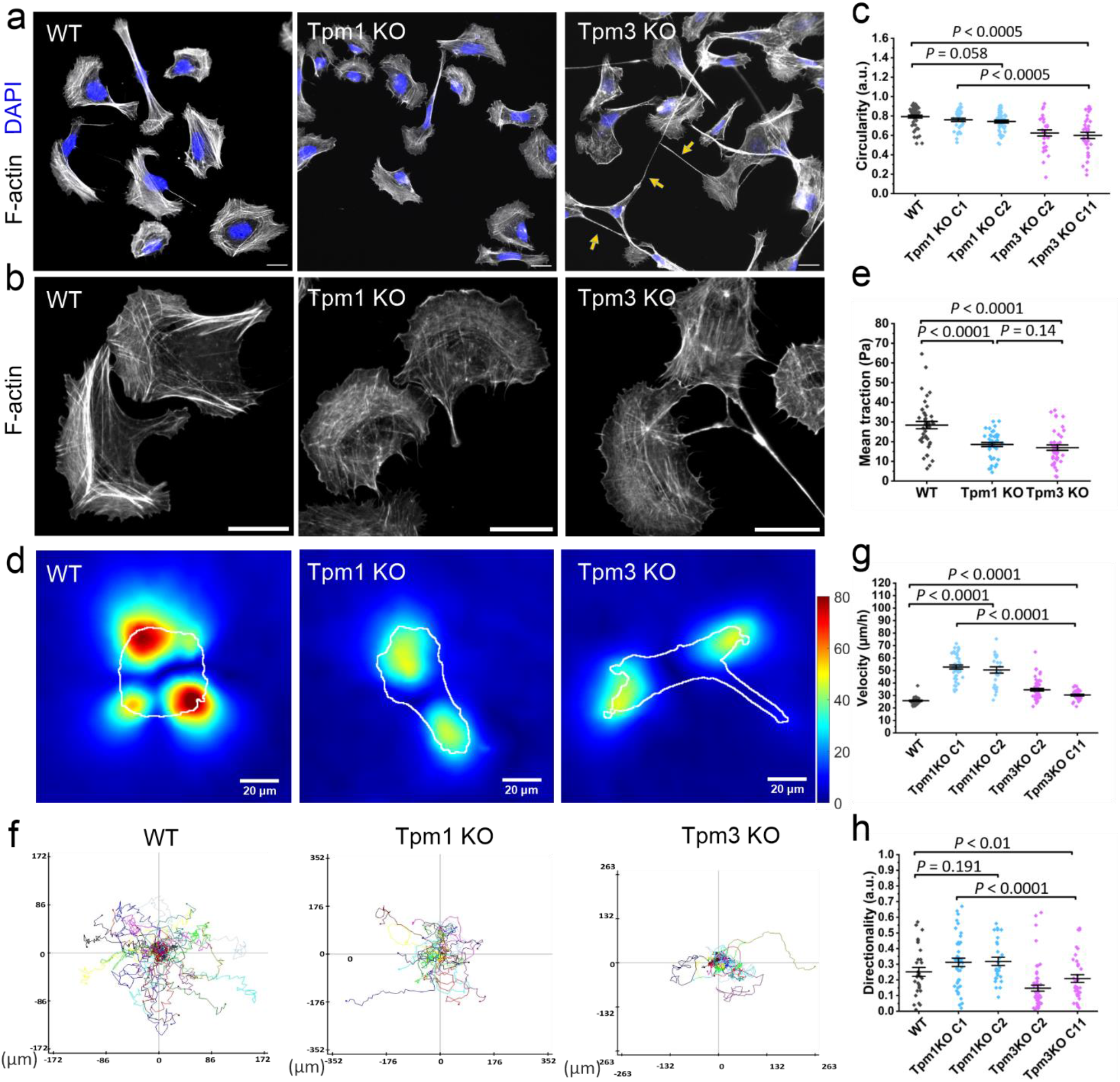
Effects of tropomyosin-1 and tropomyosin-3 depletions on cell morphology, force production, migration, and the actin cytoskeleton. (a) Representative wide-field images of wild-type, Tpm1 knockout, and Tpm3 knockout U2OS cells stained for F-actin (phalloidin) illustrating the morphological differences between the Tpm1 knockout and Tpm3 knockout cells. The arrows highlight the abnormal tails of the Tpm3 knockout cells. Scale bars, 20 μm. (b) Representative wide-field images of wild-type, Tpm1 knockout, and Tpm3 knockout cells stained for F-actin (phalloidin), demonstrating defects in the stress fiber networks of the Tpm1 knockout and Tpm3 knockout cells (c) Cell circularity analysis of wild-type (n=42), Tpm1 knockout; clone 1 (C1) (n=44) clone 2 (C2) (n=45), and Tpm3 knockout; clone 2 (C2) (n=37) clone 11 (C11)(n=38) cells after 90 min of plating. The data represents mean ± SE. (d) Representative traction force maps of wild-type, Tpm1KO and Tpm3KO cells plated on 9.6 kPa hydrogels coated with fibronectin. Scale bar 20 μm. (e) Mean traction (per cell) of wild-type (n=43), Tpm1KO (n=41) and Tpm3KO (n=42) cells on 9.6 kPa hydrogels coated with fibronectin. The data represents mean ± SE. (f) Random migration trajectory maps of wild-type, Tpm1 knockout, Tpm3 knockout cells plated on fibronectin-coated substrate. (g-h) Analysis of random migration rates and cell directionality of wild type (n=30), Tpm1 knockout; clone 1 (C1) (n=42) clone 2 (C2) (n=27), and Tpm3 knockout; clone 2 (c2) (n=55) clone 11 (C11) (n=34) cells. The data represents mean ± SE.

Interestingly, although Tpm1 and Tpm3 depletions resulted in similar effects on stress fiber networks, the knockouts affected cell morphology and migration in different ways (Fig. 3a, b, S4f). During the initial phase of cell spreading (90 minutes post plating) on a fibronectin-coated glass surface, wild-type and Tpm1 knockout cells displayed relatively round and non-polarized morphology, whereas Tpm3 knockout cells were more elongated and polarized. These morphological differences were consistent also in other Tpm1 and Tpm3 knockout clones (Fig. 3c) and were also retained after the Tpm1 and Tpm3 knockout cells were fully spread and polarized (8 hours post plating) (Fig. S4g). Interestingly, unlike wild-type and Tpm1 knockout cells, the majority of Tpm3 knockout cells exhibited abnormally long tails (Fig. 3a), suggesting that Tpm3 is critical for proper tail-retraction during cell migration. To explore this further, we examined random migration of cells plated on fibronectin. By tracking the movement of nuclei, we revealed that the Tpm1 knockout cells migrated with a higher velocity compared to Tpm3 knockout and wild-type U2OS cells (Fig. 3f, g, Supplementary movies 3, 4). On the other hand, Tpm3 knockout cells, but not Tpm1 knockout cells, displayed defects in directionality during migration (Fig. 3f, g, h, Supplementary movie 5), and exhibited defective tail retraction (Fig. 3a; Supplementary movie 5). Thus, Tpm1 and Tpm3 depletions result in opposite effects on cell morphology, polarity, and migration, indicating that the two actin filament populations, specified by Tpm1.6 and Tpm3.2, regulate these cellular processes via distinct molecular mechanisms.

### Opposite functions of Tpm1.6 and Trpm3.2 in spatio-temporal dynamics of focal adhesions

The localisation of Tpm1.6-actin and Tpm3.2-actin to different actin filament ‘layers’ in FAs (Fig. 2g), and their distinct effects on cell morphology and migration (Fig. 3) suggest that these proteins may regulate the organization or dynamics of FAs by different mechanisms. Interestingly, the density of vinculin-positive FAs was significantly increased in the Tpm3 knockout cells compared to the wild-type cells, whereas the adhesion density was unaffected in the Tpm1 knockout cells (Fig. 4a, b). Moreover, the Tpm1 knockout cells displayed almost a complete lack of large (area >3 μm^2^) adhesions, whereas the adhesion size-distribution was less affected by depletion of Tpm3 (Fig. 4d). Perhaps most interestingly, Tpm1 and Tpm3 knockouts had opposite effects on spatial distribution of adhesions in cells. In Tpm1 knockout cells, the adhesions were predominantly localized at the leading edge (~70 % of adhesions were within 5 μm distance from the leading edge), whereas the majority (>60 %) of adhesions in Tpm3 knockout cells were located at the cell centre or cell rear (Fig. 4c). Finally, Tpm1 knockout cells had a significantly higher number of active integrin-β1 positive adhesions, whereas the total active integrin-β1 intensity per cell was significantly lower in the Tpm3 knockout cells (Fig. S5a-c). These data suggest that Tpm1 limits integrin activation in nascent adhesions, thus supporting their maturation at the leading edge, whereas Tpm3 supports integrin activity in adhesions and may thus influence turnover of already existing focal adhesions.

**Figure 4.**
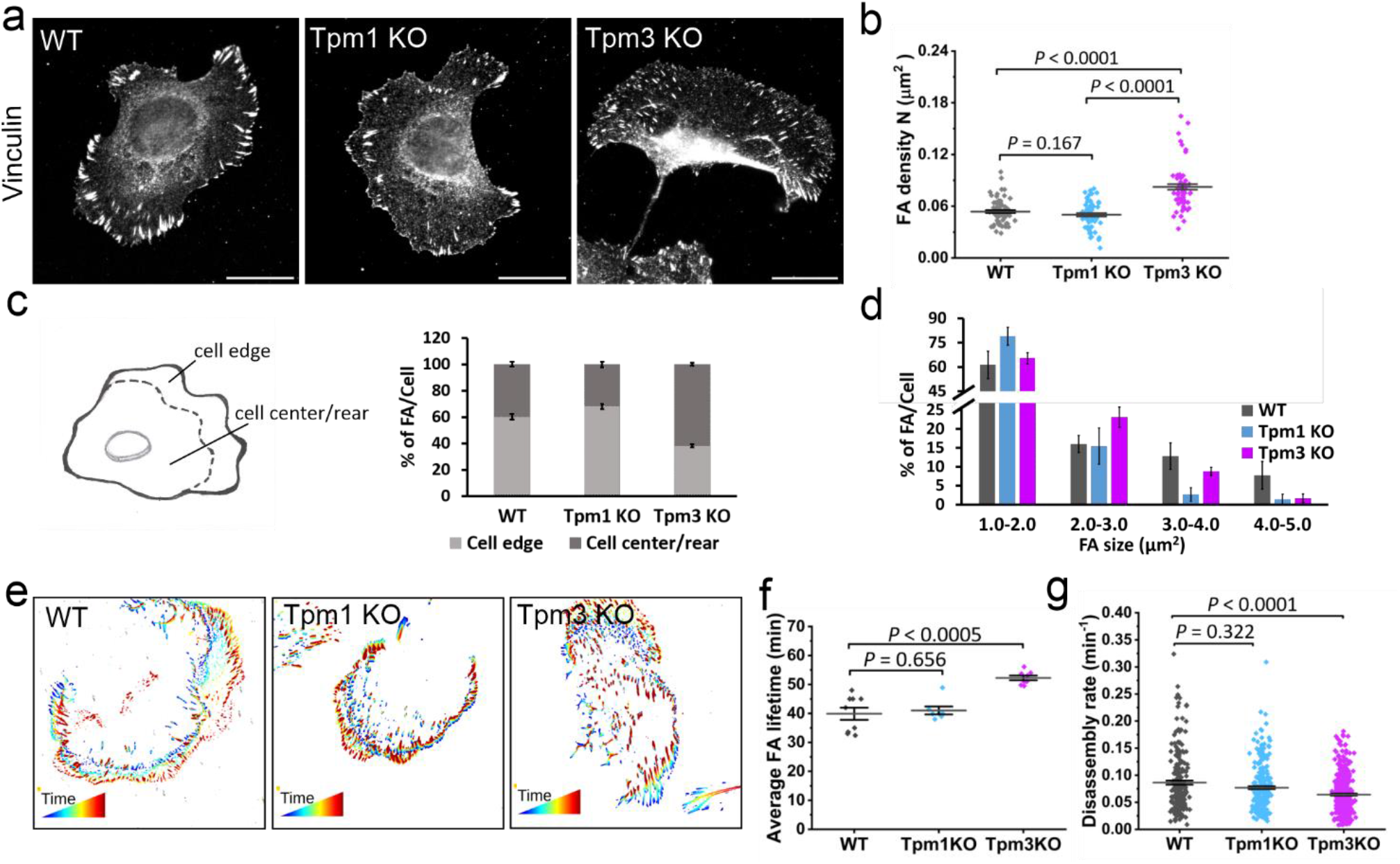
Depletions of tropomyosin-1 and tropomyosin-3 result in opposite effects on focal adhesion distribution and dynamics. (a) Representative wide-field images of wild-type, Tpm1 knockout, and Tpm3 knockout U2OS cells plated on fibronectin-coated coverslips and stained for focal adhesions (vinculin antibody). Scale bars, 20 μm. (b) Focal adhesion density analysis from wild-type (n= 70), Tpm1 knockout (n= 62), and Tpm3 knockout (n= 63) cells. The data represents mean ± SE. (c) Distributions of focal adhesions at the cell edge (within 5 μm from the leading edge) vs. cell centre/rear. Quantification of the percentage of focal adhesions located in these regions in wild-type (n= 30), Tpm1 knockout (n= 30) and Tpm3 knockout (n=25) cells are shown in the graph. (d) Quantification of the size distributions of focal adhesions in wild-type (n= 30), Tpm1 knockout (n= 30) and Tpm3 knockout (n=25) cells. The data represents mean ± SE. (e) Representative examples of temporal color-coded TIRF microscopy time-lapse movies of miRF670-Paxillin expressing cells depicting the dynamic mode of focal adhesions (generated from FAAS). (f) Quantitative analysis of average focal adhesion lifetimes from TIRF microscopy time-lapse movies of miRFP670-Paxillin expressing wild-type (n=9 cells), Tpm1 knockout (n=7 cells), and Tpm3 knockout (n=8 cells) cells. The data represents mean ± SE. (g) Analysis of focal adhesion disassembly rates calculated from TIRF microscopy time-lapse movies of miRFP670-Paxillin expressing cells by using FAAS. Wild-type (n=9 cells), Tpm1 knockout (n=7 cells), and Tpm3 knockout (n=8 cells). The data represents mean ± SE.

To determine if the differences in spatial distributions of FAs in Tpm1 and Tpm3 knockout cells are linked to altered adhesion dynamics, we performed TIRF live-cell imaging of wild-type, Tpm1 knockout, and Tpm3 knockout cells transiently expressing miRFP tagged paxillin (Fig 4e, Supplementary movies 6-8). Analysis of the obtained movies by Focal Adhesion Analysis Server (FAAS) ^50, 51^, demonstrated that the average lifetime of adhesions per cell was unaffected by the depletion of Tpm1, whereas average lifetimes of adhesions were increased by ~30% in the absence of Tpm3 (Fig. 4e-f). Analysis of the dynamics of individual paxillin-positive adhesions by FAAS also provided evidence that the FA disassembly rates were significantly diminished in the Tpm3 knockout cells, consistent with their increased adhesion lifetimes (Fig. 4g). Together, these data show that Tpm1.6-actin filament arrays, which may link adhesions to stress fibres, are important for FA maturation. On the other hand, Tpm3.2-actin filament arrays residing towards the bottom of adhesions are crucial for proper FA disassembly at the cell centre and cell rear. Consequently, depletions of Tpm1 and Tpm3 lead to opposite effects in the subcellular distribution and dynamics of FAs.

### Tpm3.2 controls focal adhesion disassembly through KANK-dependent microtubule-targeting

FA disassembly must be accurately regulated to maintain proper spatio-temporal distribution of adhesions for cell migration and morphogenesis. Microtubule targeting to focal adhesions is crucial for adhesion turnover, but the underlying molecular details are incompletely understood ^12,13^. Given the FA disassembly defects in the Tpm3 knockout cells, we examined the effects of Tpm3 depletion on the microtubule organization in interphase cells. Interestingly, compared to the wild-type and Tpm1 knockout U2OS cells, the Tpm3 knockout cells displayed microtubules with ‘criss-cross/tangled’ (disorganized) pattern. Moreover, the individual microtubules of Tpm3 knockout cells were less straight/aligned, and often bent at the cell periphery (Fig. 5a; Fig. S6a). Blind analysis of cells stained for α-tubulin revealed that whereas ~50 % of wild-type cells had predominantly aligned microtubules organized in parallel arrays, such ‘normal’ organization of microtubules was not detected in the Tpm3 knockout cells. In contrast, > 70 % of Tpm3 knockout cells exhibited severely disorganized microtubule arrays characterized by bent microtubules, which formed a ‘criss-cross’ network (Fig. 5b). In addition to defects in the general organization of the microtubule networks, both the subcellular distribution and morphology of microtubule plus-end tracking protein EB1 foci were affected by Tpm3 depletion (Fig. 5d; Fig. S6b).

**Figure 5.**
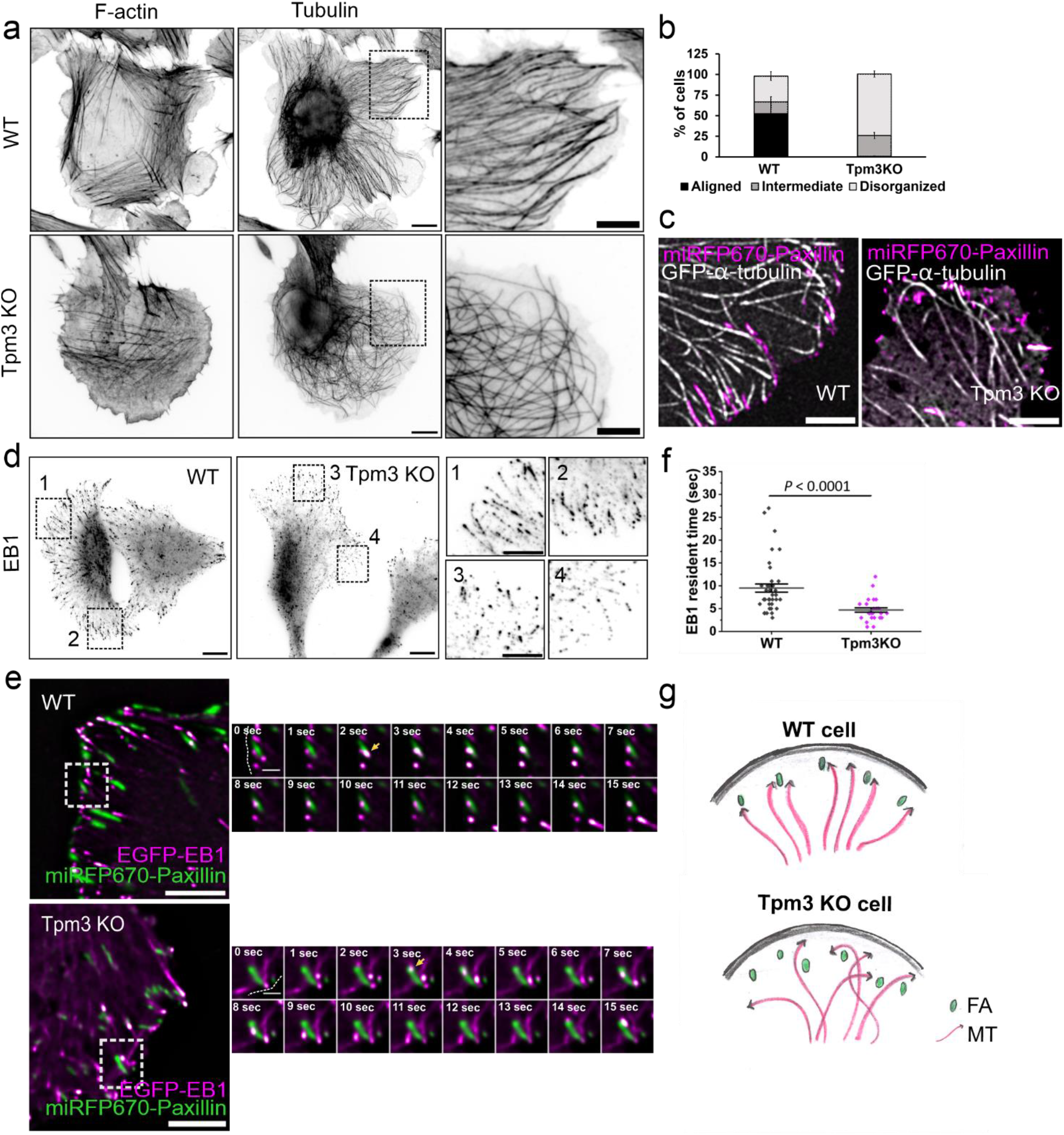
Tpm3-actin filaments are critical for proper microtubule organization and focal adhesion targeting. (a) Wide-field images of wild-type and Tpm3 knockout U2OS cells stained for F-actin (phalloidin) and microtubules (α-tubulin antibody). The panels on the right are magnified images of the regions at the cell periphery indicated with black boxes in the whole cell images. Scale bars, 10 μm and 5 μm, respectively. (b) Blind-analysis of the percentage of cells displaying aligned, intermediate and tangled microtubule networks in wild-type (n=219) and Tpm3 knockout (n=344) cells. The data are from three independent experiments and represent mean ± SE. (c) TIRF microscopy images of wild-type U2OS and Tpm3KO cells expressing miRFP670-Paxillin and GFP-α-tubulin demonstrating diminished targeting of microtubules to focal adhesions. Scale bars, 5 μm. (d) Wide-field images of wild-type and Tpm3 knockout U2OS cells stained for microtubules plus-end tracking protein EB1. The panels on the right (1-4) are magnified images of regions at the leading edge and at the cell rears, indicated with black boxes in the whole cell images. Scale bars, 10 μm 5 μm, respectively. (e) TIRF time-lapse images of wild-type and Tpm3 knockout cells expressing EGFP-EB1 and miRFP670-Paxillin. Selected time-lapse frames from the magnified areas (indicated by white boxes) are shown on the right as merged frames. Yellow arrows highlight the point of initial contact of EB1 with focal adhesions. Scale bars, 5 μm and 1 μm, respectively. (f) Analysis of the EB1 resident times in focal adhesions analysed from TIRF time-lapse movies of cells expressing EGFP-EB1 and miRFP670-Paxillin. Wild-type (n= 40 EB1 foci from 5 movies) and Tpm3 knockout (n= 26 EB1 foci from 5 movies) cells. The data represents mean ± SE. (g) Schematic representation of the organization of microtubule plus-ends in wild-type and in Tpm3 knockout cells based on the data presented here.

To examine the interplay between microtubules and FAs, we performed TIRF live-cell imaging on U2OS cells co-expressing GFP-α-tubulin and miRFP-paxillin, as well as on cells co-expressing EGFP-EB1 and miRFP-paxillin. Wild-type cells exhibited relatively stable association of microtubule plus-ends with FAs, as well as ‘poking behaviour’, where microtubules transiently targeted adhesions. In the Tpm3 knockout cells, however, microtubules were poorly targeted to FAs, and often bypassed adhesions and subsequently continued to elongate parallel to the cell edge before disassembling (Fig. 5c, supplementary movie 9). Furthermore, TIRF imaging revealed significantly shorter resident times of EB1 foci with FAs in Tpm3 knockout cells as compared to wild-type cells (Fig. 5e-g, supplementary movie 10).

To elucidate the molecular mechanism by which Tpm3.2–actin filaments mediate microtubule targeting to FAs, we focused on a multi-protein complex referred as ‘cortical-microtubule-stabilizing complex’ (CMSC), consisting of KANKs, ELKS, liprins, LL5β and CLASPs ^52^. Among the CMSC proteins, KANK directly binds to the core FA component talin. Two isoforms of this protein family, KANK1 and KANK2, localize to the rim of FAs and are critical factors in microtubule-dependent adhesion disassembly ^15, 17, 53^. TIRF imaging revealed that endogenous KANK1 and KANK2 localized either to the adhesion periphery or in the close vicinity of FAs in wild-type cells, whereas in Tpm3 knockout cells these proteins appeared to display more diffuse distributions (Fig. 6a; S7a-b; S8a). Next we investigated the effects of Tpm3 depletion on the turnover of GFP-KANK1 at miRFP-paxillin labelled FAs using FRAP. GFP-KANK1 localized to miRFP-paxillin positive adhesions in the wild-type cells, and somewhat less extensively in the Tpm3 knockout cells (Fig. 6b). Importantly, FRAP analysis revealed that GFP-KANK1 was quite stably associated with FAs in wild-type cells (with a stable fraction of ~50 %), whereas GFP-KANK1 displayed much more rapid dynamics in adhesions of Tpm3 knockout cells (with apparently no stable fraction) (Fig. 6b, c, S8b). Because KANK acts as a bridge that connects FAs to other components of CMSCs, we also examined the possible effects of Tpm3 depletion on ELKS (by antibody staining) and CLASP2 (by expressing mCherry-CLASP2). CLASP2 was enriched in the cell periphery with occasional co-localization with FAs in wild-type cells. Also, ELKS typically localized to the vicinity of FAs. However, in the Tpm3 knockout cells both CLASP2 and ELKS showed predominantly diffuse cytoplasmic localization (Fig S7c, d).

**Figure 6.**
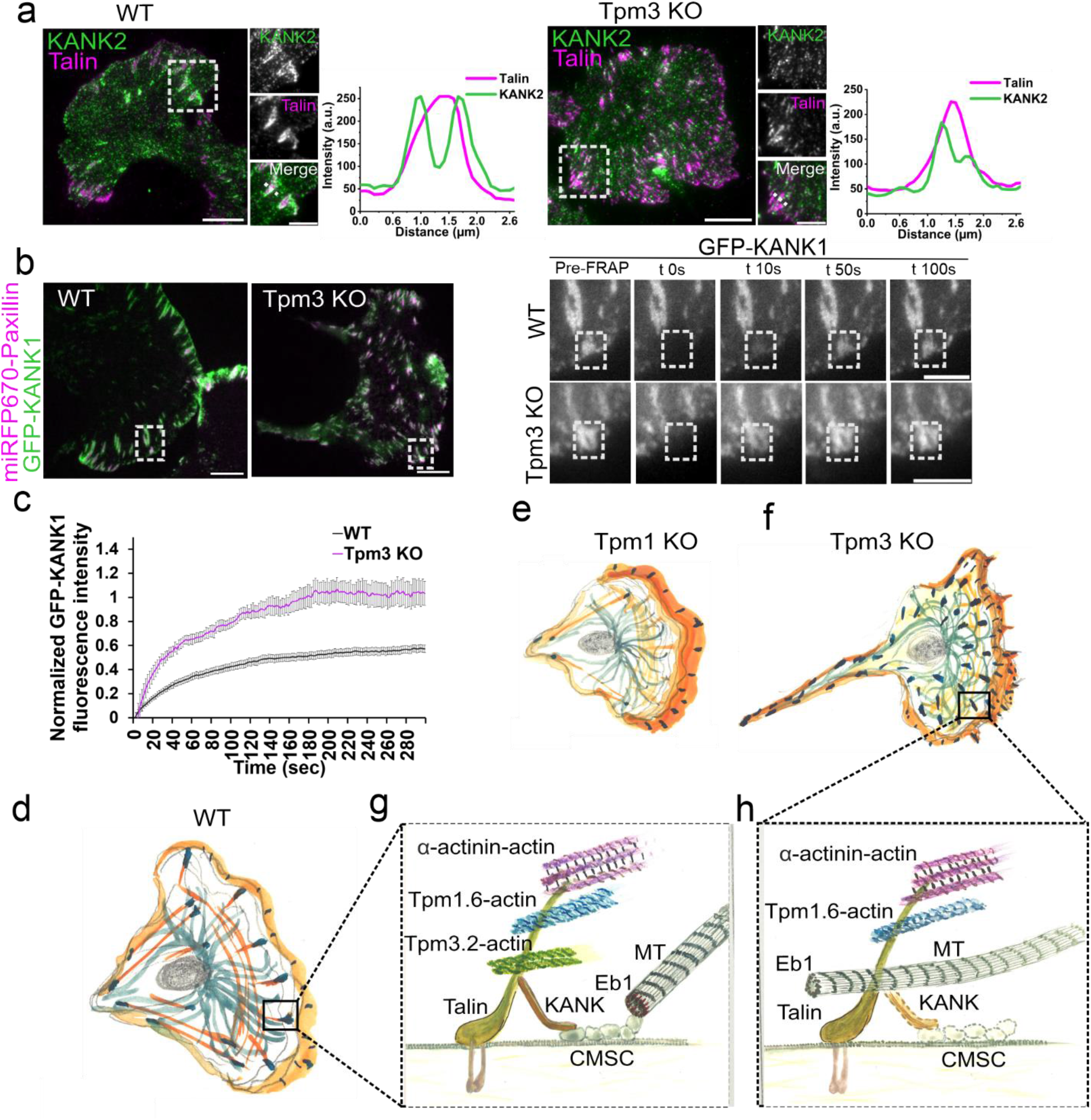
Tropomyosin-3 regulates microtubule – focal adhesion interactions by stabilizing KANK at focal adhesions. (a) TIRF images of wild-type and Tpm3 knockout cells stained for endogenous KANK2 and talin. The panels to the right are magnified images of the regions indicated with white boxes in the whole cell images. Scale bars, 10 μm and 5 μm, respectively. The intensity line scans on the right are from the selected adhesion regions demonstrating compromised localization of KANK2 at the rim of the focal adhesions in Tpm3 knockout cells. (b) Representative examples of GFP-KANK1 time-lapse images (from the FRAP experiment) from wild-type and in Tpm3 knockout cells. Scale bars, 10 μm. The panels on the right are magnified images of the regions indicated with white boxes, and represent selected time-frames of the FRAP experiments. The time-point ‘Pre’ is a frame before bleaching and ‘0 s’ is the first frame after bleaching. The white boxes indicate the bleached regions. Scale bars, 5 μm. (c) Quantification of the fluorescence recovery of GFP-KANK1 in focal adhesions of wild-type and Tpm3 knockout cells. Graph shows mean curves ± S.D. over time. The measurements are from (n=29 KANK1 patches from 7 movies) wild-type and (n=38 KANK1 patches from 6 movies) Tpm3 knockout cells. (d-f) Schematic representation of polarized wild-type, Tpm1 knockout and Tpm3 knockout cells, displaying the spatial organization of focal adhesions (black), actin (orange) and microtubule networks (green). Tpm1 depletion leads to defects in focal adhesion maturation, whereas Tpm3 depletion results in defective targeting of microtubules to focal adhesions, and accompanied problems in adhesion disassembly and tail retraction during cell migration. (g) Schematic representation of microtubule-dependent focal adhesion disassembly in wild-type cells, where Tpm3.2-actin filaments (green) close to the bottom of adhesion stabilize KANK (orange) and cortical microtubule stabilizing complex (CMSC) at adhesions. Thus, Tpm3.2-actin filaments facilitate targeting of microtubule (MT, yellow) plus-ends to the adhesion. (h) In the absence of Tpm3.2-actin filaments, the stable association of KANK with adhesions is lost, and this results in defective targeting of microtubule plus-end to the adhesion.

Together, these results reveal that Tpm3.2 is important for stabilizing KANK and its interaction partners at adhesions. Hence, Tpm3.2 is critical for proper microtubule – FA interplay, and the loss of Tpm3 consequently leads to defective microtubule-dependent disassembly of FAs.

## DISCUSSION

Our study identifies two new, spatially and functionally distinct actin filament layers in the nanoscale architecture of FAs. These layers are specified by Tpm1.6-decorated actin filaments, which localize slightly beneath the previously described ‘actin-regulatory layer’, and Tpm3.2-decorated actin filaments, which are positioned at the bottom of FA towards the previously described ‘force transduction layer’. Importantly, the two tropomyosin-actin layers have opposite roles in focal adhesion regulation: the Tpm1.6-actin layer controls integrin activation, adhesion maturation and cell migration, whereas the Tpm3.2-actin layer promotes adhesion disassembly by facilitating microtubule targeting to FAs.

Our findings of Tpm1.6 and Tpm3.2-actin layers being spatially distinct are consistent with previous biochemical studies demonstrating that Tpm1.6 and Tpm3.2 bind actin filaments in a mutually exclusive manner and hence cannot co-exist on the same actin filament. Moreover, both tropomyosins also appear to compete for actin-binding with α-actinin ^41,45^, demonstrating that Tpm1.6, Tpm3.2 and α-actinin indeed segregate to distinct actin filament arrays within FAs. Hence, the architecture of FAs is more complex than depicted in current models. This also agrees with electron microscopy studies highlighting the heterogenous organization of actin filaments within adhesions ^54–56^. In the future, it will be interesting to identify other components associating with the newly identified Tpm1.6- and Tpm3.2-actin layers within FAs, because different Tpm isoforms regulate interactions of other proteins with actin filaments to form functionally specific actin filament structures ^40^.

Our knockout studies demonstrate that the two actin filament layers, specified by Tpm1.6 and Tpm3.2, have different functions in FA dynamics. Tpm1 knockout cells display severe defects in adhesion maturation, and abnormalities in cell migration. These phonotypes were consistent in two different knockout clones, and could be rescued by expression EGFP-Tpm1.6 in the knockout cells, demonstrating that they indeed arise from the loss of Tpm1.6 (Fig. S5d, e). It is also important to note that, in addition to adhesions, Tpm1.6 localizes to stress fibers, which were thinner and less organized in the Tpm1 knockout cells compared to the wild-type cells. Thus, we propose that Tpm1.6 contributes to FA maturation by both stabilizing the stress fiber network, as well as by linking FAs to stress fibers. These functions provide a plausible explanation for the diminished traction forces, and defects in force-dependent adhesion maturation in the Tpm1 knockout cells (Fig. 6e). It is also important to note that other splice variants of the *TPM1* gene, Tpm1.8 and Tpm1.9, were earlier shown to contribute to nascent adhesion assembly. However, unlike Tpm1.6, the Tpm1.8 and Tpm1.9 isoforms do not extensively localize to FAs or to stress fibers, but instead form lamellipodial actin filament arrays, which anchor to the nascent adhesions during their assembly ^57^. Moreover, our Tpm1 knockout targeted exon 1a of the *TPM1* gene, and thus specifically depleted Tpm1.6 and closely related Tpm1.7 isoforms, without affecting Tpm1.8 or Tpm1.9 (Fig. S4a).

Whereas Tpm1.6 contributes to adhesion maturation, the Tpm3.2-actin filament layer towards the bottom of FA is important for regulated adhesion disassembly. Consequently, Tpm3 knockout cells displayed abnormal accumulation of FAs at the centre and rear (Fig. 6f). Accumulation of unusually stable adhesions also explains the abnormal morphology of the Tpm3 knockout cells, as well as defects in tail retraction during cell migration. At the mechanistic level, we revealed that microtubules fail to properly target FAs in Tpm3 knockout cells, and this results in an abnormal organization of the interphase microtubule networks characterized by ‘disorganized’ criss-crossing of microtubules, their elongation past adhesions, and growth in parallel to the cell edge. These phenotypes were consistent in the two knockout clones analysed, and could be partially rescued by expression of EGFP-Tpm3.2 in the knockout cells (Fig. S6c-e), demonstrating that these defects indeed result from the loss of Tpm3.2 protein. Interplay between Tpm3.2 and microtubules is also supported by co-immunoprecipitation work with Tpm3.1/2-specific antibody identifying tubulin, EB1, and dynein as proteins associating with Tpm3.1 or Tpm3.2 ^58^.

How do Tpm3.2-actin filaments contribute to microtubule targeting to adhesions? The diminished resident times of microtubule plus-ends with FAs in the Tpm3 knockout cells correlated with less extensive localizations of KANK1, KANK2 and ELKS around FAs. These proteins were shown to be critical for targeting microtubule plus-ends to FAs ^16^. The precise mechanism by which microtubules promote FA disassembly are still incompletely understood, but at least in some cell-types it involves local inhibition of RhoA ^17^. Whether this is also the case in U2OS cells used in our study remains to be shown, but the global levels of active RhoA were decreased, rather than increased in the Tpm3 knockout cells (Fig. S4h). In addition to local regulation of RhoA, microtubules contribute to adhesion turnover through delivery of autophagosomes and matrix metalloproteinases to FAs ^18, 19, 59^. Moreover, local acetylation of microtubules by αTAT1, which binds talin at FAs in a mechanosensitive manner, can release GEF-H1 from microtubules to enhance RhoA-mediated stress fiber contractility ^60^. Thus, future work is needed to uncover the specific roles of these processes in Tpm3.2-KANK-microtubule-dependent FA disassembly.

Although KANK proteins still accumulated to FAs to some extent in the Tpm3 knockout cells, the dynamics of KANK1 were drastically altered by Tpm3 depletion. In wild-type cells, approximately 50 % of GFP-KANK1 was stably associated with FAs, but in the absence of Tpm3 the immobile fraction was lost and the majority of the GFP-KANK1 fluorescence recovered in >30 seconds. Interestingly, the dynamics of KANK1 in Tpm3 knockout cells was similar to the previously reported FA dynamics of the isolated talin-binding KN domain of KANK1, which also was much more dynamic than full-length KANK1 and lacked a stable fraction in FAs ^53^. Therefore, stable association of KANK1 with adhesions requires both direct interaction with talin through the KN domain KANK1, as well as the presence of Tpm3.2-actin filaments (Fig. 6g-h). An earlier study also provided evidence that external force is required to stabilize the KANK-talin interaction ^53^. Thus, it is tempting to speculate that Tpm3.2-actin filaments associate either directly or indirectly with KANK to provide the required force to stabilize KANK-talin interaction at FA. In this context, it is also important to note that previous iPALM studies on pluripotent stem cells reported similar vertical localizations of KANK1 and KANK2 in FAs as Tpm3.2 in our experiments ^27^.

The opposite functions of Tpm1.6 and Tpm3.2 in FA turnover also provide a plausible explanation for the differential roles of these tropomyosin isoforms in cancer progression. Upregulation of Tpm3.1 and Tpm3.2 is detected in the majority of cancers and transformed cells, and a small molecule inhibitor of Tpm3.1 was shown to kill cancer cells in vitro and in xenotransplants ^58, 61, 62^. On the other hand, Tpm1.6 and Tpm1.7 levels are typically decreased in transformed cells ^49, 63^.

Thus, we propose that over-expression of Tpm3.2, and the closely related Tpm3.1, leads to abnormally dynamic FAs, which allow uncontrolled cell migration and invasion. Similarly, the loss of Tpm1.6 leads to defects in adhesion maturation, and accelerated and uncontrolled cell migration, hence explaining the function of Tpm1.6 as cancer suppressor.

Collectively, our study reveals unexpected complexity in the structural organization of actin at FAs and hence sets a framework for future studies on the nanoscale localization of other FA components and their interplay with the three actin filament layers of adhesions. Our work also reveals a new critical component of FAs, Tpm3.2-actin filaments, which are important for microtubule – adhesion interactions, and hence for regulated FA turnover. Finally, our study presents a striking example of three different actin filament populations, with distinct molecular functions, working in concert to control the assembly, disassembly and function of a specific actin-based cellular structure.

## METHODS

### Reagents and Antibodies

The following antibodies were used in this study. Mouse anti-vinculin (Sigma Aldrich, V9131), mouse anti-paxillin (BD Bioscience, 610569), rabbit anti-paxillin (GeneTex, GTX125891), mouse anti-active integrin-β1 (12G10; abcam, ab30394), rabbit anti-NM-IIA (BioLegend, 909801), rabbit anti-NM-IIB (BioLegend, 909901), rabbit anti-vinculin (abcam, 73412), mouse anti-α-tubulin (Digma Aldrich, T5769), mouse monoclonal anti-EB1 (1A11/4; Santa Cruz, 47704), rabbit anti-KANK1 (Bethyl laboratories, A301-882A), rabbit anti-KANK2 (Sigma-Aldrich, HPA015643), mouse anti-talin (Sigma Aldrich, T3287), tubulin, rabbit anti-ERC1 (ELKS) (Sigma-Aldrich, HPA019513), rabbit anti-Y118-phosphorylated paxillin (Cell Signaling Technology, 2541), antibody against Tropomyosin1, 2; mouse anti-Tropomyosin 1 and 2 (T2780, clone name-TM311, Sigma-Aldrich, 014M4782) and mouse anti-Tropomyosin 3 (CG3) ^47^. 4’,6’-diamidino-2-phenylindole (DAPI) (Thermo Fisher Scientific, D1306) was utilized to detect the nuclei, whereas Alexa Fluor 488- and 568-Phalloidin (Thermo Fisher Scientific, A12379/12380) were applied to visualize F-actin. Alexa Fluor 488- and 568-conjugated goat anti-mouse (Thermo Fisher Scientific, A-11001 and A-11031, respectively) and Alexa Fluor Plus 647-conjugated goat anti-rabbit (Thermo Fisher Scientific, A32733) were used as secondary antibodies. 5% BSA-TBS-Tween20 (0.02%) was used as a blocking buffer for the cells prior to staining as well as a diluent for the above mentioned antibodies. Polyclonal rabbit ab against GAPDH (Sigma-Aldrich, G9545) was used to probe equal loading in WB. Fibronectin (10 μg/ml, 11080938001, Merck) was used for surface-coating in live and fixed-cell imaging studies. Treated coverglass was mounted using ProLong^™^ Glass Antifade Mountant (P36980, Invitrogen).

### Cell culture and transfection

Human osteosarcoma (U2OS) cells (authenticated by ECACC through STR-profiling method to be the same origin as the original U2OS cell line; case number-13472) were maintained as previously described ^39, 64, 65^. Briefly, cells were cultured in 4.5 g/L glucose containing DMEM (BE12-614F, LONZA), supplemented with 10% fetal bovine serum (10500-064, GIBCO) and Pen-Strep-Glutamine solution (10378016, GIBCO) in a humidified atmosphere at 37 °C, 5% CO_2_ and 95% relative humidity. Cells were regularly tested for mycoplasma contamination using the Mycoalert^™^ Mycoplasma Detection Kit (LT07-418, LONZA). For live-cell imaging experiments, Fluorobrite DMEM (Gibco, A1896701), supplemented with 25 mM HEPES, glutamax and 10% FBS was used. Transient transfections were performed either with Xfect transfection reagent (Takara, 631318) or with Mirus Bio TransIT-2020 transfection reagent (MIR 5400) according to the manufacturers’ instructions. Transfections were carried out overnight before live-cell imaging or before fixation. For rescue experiments, cells were transfected for 24 hours. After transfection, cells were either fixed with 4% PFA in PBS for 15 min at room temperature or detached with 0.05% Trypsin-EDTA (Gibco, 15400054) and plated onto high-precision (#1.5H) 35 mm imaging dishes (Ibidi μ-dish high, 81158) or #1.5H coverslips coated with 10 μg/ml fibronectin. For live-cell imaging, cells were allowed to adhere for three hours prior to placing them in the imaging chamber.

### Generation of Tropomyosin 1 and Tropomyosin 3 knockout cell lines

Tropomyosin 1 and Tropomyosin 3 CRISPR knock-out cell lines were generated as described previously ^39, 64, 66^. Guide sequences targeting exon 1a (CTCGACAAGGAGAACGCCT) of the human *TPM1* gene and exon 1b (GAGAAGTTGAGGGAGAAAGG) of *TPM3* gene were selected based on the CRISPR Design tool with highest on target efficiency scores. Oligonucleotides for cloning guide RNA into pSpCas9 (BB)-2A-GFP vector (48138; a gift from F. Zhang, Addgene, Cambridge, MA) were designed as described previously ^63^. Transfected cells were sorted with FACSAria II (BD), using low intensity GFP-positive pass gating, as single cells onto a 96-well plate, supplemented with DMEM containing 20% FBS and 10 mM HEPES buffer. For this study, two CRISPR clones were selected for each KO based on the lack of detectable Tpm1 and Tpm3 proteins by Western blot. All the data presented in the manuscript for Tpm1 and for Tpm3 knockouts are from clones 1 and 2, respectively. Data for Tpm1 knockout clone 2 and for Tpm3 knockout clone 11 are shown in Figure 3. For further validation of tropomyosin knockouts we performed Sanger Sequencing (Eurofins genomics sequencing service) and next generation sequencing (NGS) (Illumina MiSeq.). For Tpm1 knockout clone 1, CRISPR resulted in deletion of 17 nucleotides that eventually leads to a frameshift and appearance of STOP codon soon after the deletion site. The results were confirmed by NGS (MiSeq, Illumina). For Tpm3 knockout clone 2, CRISPR resulted in insertion of a single nucleotide (A) that eventually leads to a frameshift and appearance of STOP codon soon after the deletion site. NGS was performed at the DNA Sequencing and Genomics Laboratory (BIDGEN) laboratory (Institute of Biotechnology, University of Helsinki, Finland).

### Generation of EGFP-CAAX cell line

The pEGFP-CAAX plasmid was made by adding the CAAX-motif (encoding the peptide CMSCKCVLS) of H-Ras to the C-terminus of EGFP by using inverse PCR. EGFP-CAAX was then cloned into the safe harbor vector pSH-FIRE-P-AtAFB2 (Addgene, #129715) that was integrated into the safe harbor of U2OS cells using CRISPR/Cas9. Stable cells expressing EGFP-CAAX were obtained by puromycin selection.

### Immunofluorescence staining

Cells were cultured on 10 μg/mL fibronectin-coated (Sigma-Aldrich, L2020) coverslips or on 35 mm imaging dishes (Ibidi μ-dish high). Except for microtubule and EB1 staining, cells were fixed with 4% PFA in PBS at RT for 15 mins, washed several times with PBS, permeabilized with 0.2% Triton X-100 in PBS at RT for 7 mins and washed again with PBS. Cells were blocked with 5% BSA in PBS for 1 hour at RT and incubated with primary antibodies in 5% BSA for 1 h at room temperature. Cells were washed several times with PBS-T (0.02% Tween-20 in 1X PBS), and incubated with secondary antibodies for 1 hour at RT. After several washing with PBS-T, coverslips were mounted onto microscopes slides with ProLong^™^ Glass Antifade Mountant (P36980, Invitrogen). Microtubule staining was performed as described before ^68^. In brief, cells were fixed with 3% PFA + 0.025% Glutaraldehyde in Cytoskeletal Buffer (CB) supplemented with 10% Sucrose (Sucrose was added to the buffer just before the use) stock solution containing 10 mM HEPES (H3375, Sigma) at pH 6.1, 138 mM KCl (P3911, Sigma), 3 mM MgCl2 (208337, Sigma) and 2 mM EGTA (E3889, Sigma) at room temperature for 10 mins. Cells were washed with PBS and incubated with a solution of 1 mg/mL sodium borohydride in PBS for 10 mins at RT. Cells were then permeabilized with 0.2% Triton X-100 in PBS at RT for 7 mins. For EB1 staining, cells were fixed with cold methanol for 5 mins on ice.

### Western Blotting

Cell lysates were prepared by washing the cells once with cold PBS and scraping them into lysis buffer (50 mM Tris-HCl pH 7.5 150 mM NaCl, 1 mM EDTA, 10% Glycerol, 1% Triton X-100) supplemented with 1 mM PMSF, 10 mM DTT, 40 mg/ml DNase I and 1 mg/ml of leupeptin, pepstatin, and aprotinin. All preparations were conducted on ice. Protein concentrations were determined with Bradford reagent (#500-0006, Bio-Rad) and equal amounts of the total cell lysates were mixed with Laemmli Sample Buffer, boiled, and run on 4%–20% gradient SDS-PAGE gels (#4561096, Bio-Rad). Proteins were transferred to nitrocellulose membranes with Trans-Blot Turbo transfer system (Bio-Rad) using Mini TGX gel transfer protocol. Membranes were blocked in either 5% milk-TBS with 0.1% Tween20 (TBS-T) or with 5% BSA for one hour at RT. Primary and secondary antibodies were diluted into fresh blocking buffer for overnight incubation at 4°C and one hour at room temperature, respectively. Protein detection from the membranes was performed with Western Lightning ECL Pro substrate (PerkinElmer NEL121001EA).

## MICROSCOPY

### Widefield

Widefield imaging for IF samples were performed with Leica DM6000B wide-field fluorescence microscope equipped with a 63x/1.40-0.60 HCX PL APO Lbd.bl. Oil wd=0.10 and 40x/1.25-0.75 HCX PL APO Oil wd=0.10. The images were acquired using Hamamatsu Orca-Flash4.0 V2 sCMOS camera with image resolution 2048 × 2048 pixels.

### TIRF

Fixed cell imaging on 35 mm imaging dishes (Ibidi μ-dish high) in 1× PBS was performed with Ring-TIRF module of Deltavision OMX SR (Cytiva) with 60×/1.49NA Apo N oil objective (Olympus), at RT, using 1.518 RI immersion oil and 488, 561 and 607 nm diode lasers. 5×5 FOVs (1024 x 1024) including 10% overlap, were captured manually, followed by moving to a new area, at least over six times the FOV to another direction, at random positions on the imaging dish. Live cell imaging was performed as fixed cell TIRF experiments, but with following exceptions: imaging was performed with 1.522 RI immersion oil and imaging chamber with controlled humidified atmosphere of 5% CO_2_ and 37°C was utilized. Sample illumination with 488, 561, and 607 nm diode lasers was detected and recorded with three respective sCMOS cameras and controlled through Acquire SR 4.4 acquisition software. The captured 1024 × 1024 time-lapse videos had a pixel size of 0.08 μm (x/y). The time-lapse imaging for miRFP670-paxillin expressing cells was performed with 30 s interval for a total of 2 hours. 607 laser was used at 25% laser power with 100 ms exposure time. The time-lapse imaging for cells co-expressing miRFP670-paxillin either with GFP-α-tubulin or EGFP-EB1 was performed with 3 s and 1 s intervals for a total of 5 mins and 1 min, respectively. 488 and 607 lasers were used at 15 % laser power with 50 ms exposure time. Obtained time-lapse series were deconvolved and channels aligned with SoftWoRx 7.0. Prior the onset of live-cell imaging, cells were allowed to settle within the imaging chamber for 30 mins. FRAP experiments were also performed with Deltavision OMX SR. Live-cell imaging for tropomyosin recruitment to adhesions was performed with TIRF module of ONI Nanoimager S with 100x/1.49 NA oil objective at 37°C on 35 mm imaging dishes (Ibidi μ-dish high). Sample illumination with 488, 561, 640 nm lasers was detected and recorded with sCmos camera with field of view of 50μm x 80μm. 488, 561 and 640 lasers were used at 3% laser power with a 400 ms exposure time. The time-lapse imaging for miRFP670-Paxillin, EGFP-Tpm1.6, mRuby2-Tpm3.2 expressing cells was performed with 30 s interval for 1.5 hrs.

### iPALM

iPALM imaging was performed similar to as previously described ^3, 27^ on U2OS cells plated on 25 mm diameter round coverslips containing gold nanorod fiducial markers (Nanopartz, Inc.), passivated with a ca. 50 nm layer of SiO2, deposited using a Denton Explorer vacuum evaporator, except the coverslips were not coated with any ECM protein for plating the cells. Cells were plated for 18 hrs before fixation. After fixation an 18 mm coverslip was adhered to the top of the sample and placed in the iPALM. Eos-tagged samples were excited using 561 nm laser excitation at ca. 1–2 kW/cm2 intensity in TIRF conditions. Photo-conversion of Eos was performed using 405 nm laser illumination at 2–10 W/cm2 intensity. 50,000–80,000 images were acquired at 50 ms exposure, and processed/ localised using the PeakSelector software (Janelia Research Campus; ^69^. Alexa Fluor 647-labelled paxillin was imaged similarly, but with 2–3 kW/cm2 intensity 647 nm laser excitation, and 30–40 ms exposure time in STORM-buffer containing TRIS- buffered glucose, glucose oxidase, catalase, and mercaptoethanol amine41. iPALM data were analysed using iPALM plotter (AIC, Janelia Research Campus; https://github.com/aicjanelia/ipalmplotter) and images were rendered using the PeakSelector software (Janelia Research Campus). iPALM localization data records both the fluorescent molecules localized within the FA, as well as molecules in the cytoplasmic fraction. To quantify the spatial distribution of the proteins within individual FA, we created FA mask based on the paxillin image by using MATLAB code. To render iPALM images, a single colour scheme was used from red to purple, covering the z range 0 - 250 nm, where features within FA are seen. The same colour scheme was also used for side-view (xz) images and for covering the z range 0-500 nm. For analysis of protein distributions in FA:Zcentre calculation, the three-dimensional molecular coordinates for each region (individual FA) were analysed to obtain histograms of vertical positions with 1-nm bins. The centre vertical positions (Zcentre) was determined from a Gaussian fit to the FA molecule peak. For proteins like and α-actinin, where dual peaks were observed, the fitting was done using the sum of two-Gaussian distributions with independent centre vertical position and width. After the histograms for all images and individual FA were obtained, they were combined into a single average Zcentre. Please note that the observed z positioning of talin-N and α-actinin is slightly (talin-N = ~ 80 nm; α-actinin N = 120 nm) higher than its previously reported z-position (talin-N Z_centre_ = ~55 nm; α-actinin N = 103 nm) in U2OS and endothelial cells ^3^.

### Fluorescence recovery after photo bleaching (FRAP)

The FRAP experiments were performed using a Delta Vision OMX SR microscope with a 1.3 Silicone Plan APO ×60 wide-field objective. Acquisition was performed using AcquireSR (Cytiva). A488 and A607 lasers were used at 25% and 35% laser power, respectively, with a 50 ms exposure time. Imaging was performed with 2 seconds time-lapse intervals for the total duration of 5 mins. GFP-KANK1, EGFP-Tpm1.6 or EGFP-Tpm3.2 at the adhesion were bleached with single 0.05 s pulse of A488 laser at 25% laser power

### Confocal

Confocal imaging was performed using a 3i Marianas CSU-W1 spinning disk confocal microscope. Fixed cell imaging was performed using a 100x Zeiss Plan-Apochromat 1.4 NA objective, with sample illumination with 405, 488, 561 and 640 nm lasers detected by a Hamamatsu sCMOS Orca Flash4.0 camera (2048 x 2048 pixels). For TFM, imaging was performed at 37°C using a 40x Zeiss LD C-Apochromat 1.1 NA aperture, and sample illumination with 405, 488, 561 and 640 nm lasers recorded by a Photometrics Evolve 10 mHz Back illuminated EMCCD camera (512 x 512 pixels). Due to the bright intensity of the beads used for TFM, the 561 nm laser was used at 10 % power.

### Traction force microscopy

#### TFM gel preparation and surface activation

First, 35 mm glass bottom dishes (D35-14-1, Cellvis) were incubated with 1 ml bind silane solution (7.14% Plus One Bind silane (GE17-1330-01, Sigma), 7.14% acetic acid in absolute ethanol) for 1 hour at room temperature, before being washed twice with 2 ml absolute ethanol and let to air-dry. To prepare 9.6 kPa hydrogels, 1.7 μl of sonicated (30 s on, 30 s off for 7 min) fluorescent beads (FluoSpheres^™^ 200 nm red, F8810, Life Technologies) were added to a 500 μl hydrogel mixture containing 94 μl 40% acrylamide (A4058, Sigma), 50 μl *N*,*N*’-methylenebisacrylamide solution (M1533, Sigma) in PBS and briefly vortexed. Hydrogel polymerization was induced through addition of 5 μl 10% ammonium persulfate (1610700, Bio-Rad) and 1 μl *N*,*N*,*N*’*N*’-tetramethylethylenediamine (T9281, Sigma), and the mixture was rapidly vortexed before 11.8 μl was added to the prepared dishes and a clean 13 mm glass coverslip placed on top. Hydrogels were allowed to polymerize for 1 hour at RT, before PBS was added and the coverslip was removed. To activate the hydrogel surface, gels were incubated with 500 μl 0.2 mg/ml Sulfo-SANPAH (803332, Sigma), 2 mg/ml *N*-(3-Dimethylaminopropyl)-*N*’-ethylcarbodiimide hydrochloride (03450, Sigma) in 50 mM HEPES for 30 min at RT with gentle agitation before being irradiated with UV light for 10 min. Gels were washed four times with sterile PBS before being coated with 10 μg/ml fibronectin at 4°C overnight.

#### Traction force microscopy

Cells were plated onto the 9.6 kPa gels for 4 hours prior to imaging. One hour prior to imaging, the cell media was replaced with media containing 50 mM HEPES (H0887, Sigma), 60 pM SiR-Actin (SC001, Spirochrome), 5 μg/ml Hoechst 33342 (H3570, Invitrogen) to allow visualisation of cells. To enable detection of traction forces exerted by the cells, beads were imaged before and after cell removal (using 20 μl pre-warmed 20 % SDS in milli-Q H2O). To correct for drift, pre- and post-cell removal bead images were aligned using the NanoJ-Core plugin ^66^. Bead tracking and force measurements were performed in MATLAB (Mathworks, version 2020a) using TFM software ^71^. For displacement field calculation, high resolution subsampling of beads was used, with no outward deformation expected, subpixel correlation via image interpolation selected, and a template size of 21 pixels with a maximum displacement of 20 pixels. For displacement field correction, vector outliers were filtered and a threshold for the normalized displacement residual of 2. Force field calculation was performed using Fourier transform traction cytometry (FTTC) with a regularization parameter of 0.0001. Actin cell masks were generated from sirAct images in Fiji/ImageJ and overlaid onto traction maps in R (R Core Team, 2022 (R Core Team (2022). R: A language and environment for statistical computing. R Foundation for Statistical Computing, Vienna, Austria. https://www.R-project.org/.) to obtain mean traction (Pa) per cell.

### Random cell migration assay

Phase contrast time-lapse imaging of migrating cells was conducted in continuous cell culturing platform Cell-IQ (CM Technologies). Twelve-well plates (Greiner) were coated with 10 μl/ml of fibronectin and cells were allowed to adhere for 2 hours prior to imaging. Cells were once washed with PBS and replaced with DMEM containing 10 mM HEPES prior to starting the imaging. The plate lid was switched to Cell-Secure (CM Technologies) enabling insertion of CO_2_ input and output valves. 5% CO_2_-flow was set cycling between 8 min on, 20 min off. Average migration velocity of wild-type, Tpm1 and Tpm3 knockout cells was quantified by tracking the nucleus movement in between 8 min imaging cycles for 25 hours with Cell-IQ analyzer (CM Technologies). The cells that did not collide with one another were selected for the analysis.

#### RhoA activity assay

The active RhoA was determined by using G protein-linked (G-Lisa) assay (Cytoskeleton, BK124) as described before ^72^. Briefly, cells were washed on ice with cold PBS and homogenized to ice-cold lysis buffer. Protein concentration was measured and adjusted to 0.9 mg/ml, and samples were snap-freezed with liquid nitrogen. Triplicate assays were done, active RhoA was measured according to the manufacturer’s instructions with absorbance of 490 nm.

## IMAGE ANALYSIS

### Cell morphology analysis

Cells were plated on fibronectin-coated coverslips, fixed after 90 mins and 8 hours post-plating and stained for actin (phalloidin) and nucleus (DAPI). The images were acquired either with Floid wide-field microscope equipped with Plan Fluorite 20x/0,45 objective and Sony 1.3MP 1/3” ICX445 EXview HAD CCD camera using phase transmitted light channel or with Leica DM6000B wide-field fluorescence microscope. The cell circularity and aspect ratio were quantified by using FIJI/ImageJ. Cell boundaries were drawn manually by using the free hand tool and the measurements taken using the ROI manager tool.

### Quantification of focal adhesion density

The FA density per cell was calculated according to equation described in ^60^:

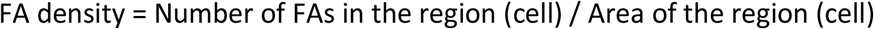

### Quantification of Tpm recruitment to focal adhesions

The dynamics of Tpm recruitment to FAs were measured using Fiji/imageJ. Visual analysis of movies identified the region and the frame where a FA started to form (paxillin signal appeared). Using a custom macro, the fluorescence signal from the all channels were measured for 10 frames before and 20 frames after the FA appearance.

### Quantification of focal adhesion spatial distribution

The FA percentage was analyzed in two regions of cells. Cell edge was defined by distance of 5 μm taken from cell periphery and rest of the region as cell center/rear. The percentage of adhesions in these regions was correlated to the total number of adhesions in the same cell.

### Analysis of focal adhesion size distribution

FA areas were quantified with Fiji/ImageJ, measuring the size of each individual adhesion with the ROI manager and freehand tool. The images were processed with the rolling ball background subtraction using 50-pixel ball radius in Fiji/ImageJ, converted to binary, and each individual adhesion analyzed with the ‘analyze particle tool’ from the adhesions’ mask created from binary images. The size threshold was set from the range of 0.08 −15 μm^2^. The value 0.08 μm^2^ was selected because our vinculin antibody visualized also unspecific background particles below the size of 0.08 μm^2^. Cells that were in contact with neighboring cells were discarded from the analysis. Adhesions were classified into four groups based on size, and the percentage ratio of the FAs in each group was obtained by dividing the FA number of individual size groups with the total number of focal adhesions in the cell.

### Quantification of focal adhesion properties

Cells transfected with miRFP670-Paxillin were allowed to adhere on fibronectin coated glass-bottom dishes (Ibidi μ-dish high) for 3 hours prior to imaging. Time-lapse images were acquired in the interval of 30 sec for the duration of 1.5 hours at TIRF plane using of Deltavision OMX SR (Cytiva) with 60×/1.49NA Apo N oil objective (Olympus). The time-lapse movies were stabilized by using the ‘image stabilizer’ plugin in Fiji/ImageJ. The movies were further processed using the Focal adhesion analysis server (https://faas.bme.unc.edu); ^50, 73^to analyze the lifetime, assembly, and disassembly rates of all paxillin-positive adhesions per cell.

### 12G10 focal adhesion analysis

Cells were plated onto fibronectin-coated polymer dishes (Ibidi, 8 well μ-slide polymer) for 3 hours before being fixed and stained for active integrin-β1 (12G10), paxillin, actin (phalloidin), and DAPI. Images were acquired using a spinning disc confocal microscope. Cell masks were generated from actin staining, and active integrin-β1 intensity calculated from the mean integrated density of the cell mask. Quantification of active integrin-β1 FAs was performed in Fiji/ImageJ using an analysis pipeline adjusted from Horzum, Ozdil and Pesen-Okvur, 2014. 12G10 images were processed using subtract background with a rolling ball radius of 20, the local contrast enhanced using the CLAHE plugin (block size = 19, histogram = 256, maximum slope = 5, no mask) and the background minimised further using EXP before thresholding to generate a binary mask. An adjustable watershed was applied to aid the detection of individual adhesions, and adhesions were analysed using the Analyze Particles tool, with a size threshold set to 0.1 to 15 μm^2^.

### FRAP analysis

FRAP data in all Figurs were analysed similar to the previously described approach ^64, 66, 74^. In brief, first the background subtraction and correction of fluorescence bleaching during imaging (the fluorescence within the region of interest was divided by that of an identical-sized region within the cell) was performed with the help of Fiji/ImageJ. Recovery rate was normalized to the prebleach intensity in every experiment. Data were analysed using Microsoft Excel.

### Quantification of microtubule organization

Cells were plated on fibronectin-coated coverslips for 8 hours before fixation and stained with tubulin antibody. The images were acquired by using a wide-field microscope. For the blind phenotypic analysis, the microtubule network was categorized into three groups based on their organization at the cell periphery: 1. ‘aligned’, 2. ‘intermediate’, 3. ‘disorganized’. The ‘intermediate’ organization was defined as where microtubule showed somewhat tangled or bended organization that was in between normal ‘aligned’ and abnormal ‘disorganized’ organization.

### Quantification of EB1 foci resident time on adhesions

Cells transfected with EGFP-EB1 and miRFP670-Paxillin were imaged at 1 sec per frame for the duration of 1 min at TIRF plane using of Deltavision OMX SR (Cytiva) with 60×/1.49NA Apo N oil objective (Olympus). The channels for the obtained time-lapse series were aligned with SoftWoRx 7.0. The obtained time-lapse series were processed using FIJI/ImageJ. Different areas at the cell periphery were selected by using the ROI tool. The obtained time-lapse series of EB1 and Paxillin were processed by using the ‘manual tracking plugin’ in ImageJ. The resident time was calculated by tracking the number of frames between the first and last frame where an individual EB1 foci localized with a paxillin positive adhesion. The first frame is defined by the event where an individual EB1 foci comes in contact with an adhesion. Thereafter, all the consecutive frames were tracked where the particular EB1 foci stays stably associated with the adhesion until the last frame before it no longer localizes with the same adhesion. The individual EB1 foci were assigned to a specific track number from the start to end of the tracking. We specifically analyzed the events where EB1 foci co-localize with paxillin, whereas the events where EB1 foci seemed to slide off on the paxillin site or were present in the close proximity of paxillin site were not considered for the resident time analysis.

### Statistics

The statistical analysis and graph construction for intensity line profile (Fig. 1b, f, S1a, b, c), bar graph generation (Fig. 4c, d, e, 5b, e) and FRAP data analysis and graph generation were performed with Excel (Microsoft). All the remaining graphs were constructed with OriginPro 2022b (OriginLab Corp.). Statistical tests were performed using two-tailed Student’s *t*-tests (Fig. 5e, S6b, c, e, Fig. 4b, g, S5b-e, Fig. 3e, S4f), One-Way ANOVA followed by Tukey’s multiple comparison’s test (Fig. 3c, g, h) and with two-tailed Mann–Whitney rank-sum tests (Sup Fig. 4g, h, Fig. 4h, Sup 6f) with OriginPro 2022b (OriginLab Corp.).

## DATA AVAILABILITY

The data supporting the findings of the study are available in the manuscript and supplementary information. Other raw data generated in the study are available from the corresponding author on reasonable request.

## ACKNOWLEDGMENTS

We thank the Biomedicum Imaging Unit (BIU) of the University of Helsinki, sponsored by HiLIFE and Biocenter Finland, and the Institute of Biotechnology Light Microscopy Unit (LMU) for technical advice and support with microscopy. Confocal microscopy was performed at the Cell Imaging and Cytometry Core, Turku Bioscience Centre, Turku, Finland, with the support of Biocenter Finland. We thank Konstantin Kogan for assistance with NGS data analysis and assembly of the related Figure. We thank Mirva Tirkkonen, Petra Laasola and Jenni Siivonen for excellent technical assistance. We also thank Aleksi Isomursu and Guillaume Jacquemet (University of Turku) for helpful discussions regarding TFM, Manuel Théry (University of Paris) for advice and discussion regarding microtubule experiments, and Juha Saarikangas (University of Helsinki) for critical reading of the manuscript. This work is supported by grants from Academy of Finland (Center of Excellence grants 346133 and 346131 to P.L. and J.I, respectively, a research fellowship 343239 to M.C. and InFLAMES Flagship Programme 337530 to J.I.), Sigrid Juselius Foundation (4708344 to P.L.), Finnish Cancer Institute (K. Albin Johansson Professorship to J.I., Agence Nationale de la Recherche (ANR-DFG JA-3038/2-1) (to R.P.), ARC (DP160101623), the Australian NHMRC (APP1100202, APP1079866) and The Kid’s Cancer Project (to P.G.).

## AUTHOR CONTRIBUTIONS

P.L., R.K. and E.K. crafted the original idea and initiated the study. R.K., R.P., T-L.C., J.I., and P.L., designed the research. R.K., K.V., M.C., J.P., J.A., E.K. carried out the experiments and analyzed the data. L.A-S., and P.G. provided reagents and expertise. R.K. and P.L. wrote the manuscript with input from all authors.

## CORRESPONDING AUTHOR

Correspondence to Pekka Lappalainen.

**Supplementary Information is available for this paper.**

